# An active light signalling pathway is necessary for ABA-induced inhibition of hypocotyl elongation

**DOI:** 10.1101/2024.01.20.576397

**Authors:** Esther Cañibano, Daniela Soto-Gomez, Juan Carlos Oliveros, Clara Bourbousse, Sandra Fonseca

## Abstract

Driven by cell elongation, hypocotyl growth is tightly controlled by light and responds to external stimuli and endogenous hormonal pathways. Hypocotyls are known to be responsive to the stress signalling hormone abscisic acid (ABA) which effectively inhibits cell elongation, but how this regulation is connected to light responses and other endogenous hormonal pathways has been a subject of limited studies. Here, we show that whereas hypocotyl elongation is sensitive to ABA in light-grown seedlings, the hypocotyl of dark-grown etiolated seedlings is ABA-insensitive. In the dark, hypocotyl sensitivity to ABA is restored in the constitutive photomorphogenic *pifq* and *cop1-4* mutants, suggesting that an active light signalling pathway is necessary for hypocotyl responsiveness to ABA. However, etiolated hypocotyls retain ABA responsiveness, as could be detected by the induction of *ABI1* and *RD29B* transcripts in response to exogenous ABA, suggesting that inhibition of hypocotyl elongation mediated by ABA does not follows the canonical ABA signalling dependent on transcription. Here, using RNA-seq analysis we identified a number of ABA differentially expressed genes (DEGs) that correlate with ABA inhibition of hypocotyl elongation, specifically in dark-grown *pifq* or light-grown WT plants, and whose expression remains unchanged by ABA treatment in dark-grown WT plants. Among these DEGs we identified a number of genes playing a role in cell elongation directly at the level of the plasma membrane, as SAURs, ion transporters, auxin flux regulators, channels, and cell wall modification enzymes. The use of the auxin transport inhibitor, NPA, revealed that in the light auxin transport impairment renders hypocotyls insensitive to ABA in WT and *pifq* plants. Thus, in the light, hypocotyl responsiveness to ABA is dependent on auxin transport and independent of PIFs. In the dark, PIFs render hypocotyls insensitive to ABA, perhaps by regulating the expression of a number of ABA DEGs, a mechanism that could allow plants to prioritize the elongation towards light, avoiding to slow-down soil emergence that could be induced by ABA signalling in case of sudden reduction of soil moisture.

## Introduction

During underground germination, seed plants like Arabidopsis have a short time window to reach the soil surface, at which point they perceive light and can become autotrophic. In this soil-buried phase, the young seedling fed by the seed reserves rapidly elongates its hypocotyl in search for light. Thus, this growth phase, driven by cell elongation, is primordial for seedling establishment and survival (Von Arnim and Deng, 1996; Gendreau *et al*., 1997). Upon light perception, hypocotyl elongation is reduced, the small cotyledons expand, turn green and photosynthetic competency is established. Under this photomorphogenic developmental program the plant grows and develops with full competencies to perceive, transduce and respond to abiotic and biotic signals that allow adaptation to the environment (Kami *et al*., 2010).

Light being a major repressor of hypocotyl elongation, mutations in light photoreceptors as cryptochromes (CRYs) or phytochromes (phy) result in seedlings with long hypocotyl phenotypes, similarly to mutants in the transcription factors (TFs) that are important for the initiation of photomorphogenic responses as ELONGATED HYPOCOTYL 5 (HY5) or HY5 HOMOLOG (HYH) (Chory, 1992; Oyama *et al*., 1997; Holm *et al*., 2002). Once photoreceptors are activated by light they render the CONSTITUTIVE PHOTOMORPHOGENESIS 1 (COP1) - SUPPRESSOR OF PHYA-105 (SPA) complexes ineffective to bind to and to promote the degradation of the target transcription factors (Liu *et al*., 2011; Zuo *et al*., 2011; Lu *et al*., 2015; Sheerin *et al*., 2015; Ponnu *et al*., 2019). Thus, in light, HY5 and HYH accumulate and trigger a photomorphogenic phenotype typical from light grown plants. HY5 over-accumulation can also trigger constitutive stress responses mimicking high light stimulus and activating genes involved in several other stresses, suggesting HY5 is a central hub for stress responses (Osterlund *et al*., 2000; Cañibano *et al*., 2021).

In the dark, HY5 is being degraded by the COP1-SPA complex (Saijo *et al*., 2003; Zhu *et al*., 2008). Independently of light conditions COP1 itself is recycled and targeted for degradation by the DET1 containing E3 ubiquitin ligase complex (Osterlund *et al*., 2000; Cañibano *et al*., 2021). Thus, impairment in COP1 or DET1 functions results in short hypocotyls and in deetiolated phenotypes in the dark, compatible with an active HY5 function (Chory *et al*., 1989; Deng *et al*., 1991).

The bHLH-type transcription factors PHYTOCHROME-INTERACTING FACTORS (PIFs) are active playing a central role in the dark. In this condition PIFs are released from the light activated phytochrome inhibition and are active to promote hypocotyl elongation. As so, the quadruple *pif* mutant (*pifq*), lacking PIF1, PIF3, PIF4, and PIF5 displays, in the dark, short hypocotyls and expanded cotyledons, characteristic of a photomorphogenic development (Toledo-Ortiz *et al*., 2003; Duek and Fankhauser, 2005; Castillon *et al*., 2007; Shin *et al*., 2007; Leivar *et al*., 2008; Leivar and Quail, 2011; Pfeiffer *et al*., 2014). Moreover, PIFs play a role adjusting hypocotyl growth to the length of day-night cycles, shade avoidance responses, warm temperature as well as hormones as gibberellins (GAs) and brassinosteroids (BRs). GA promotes hypocotyl elongation by triggering the degradation of DELLA repressors, releasing PIF3 and PIF4 to bind DNA (De Lucas *et al*., 2008; Feng *et al*., 2008). BR-induced hypocotyl elongation is mediated by BRASSINAZOLE-RESISTANT 1 (BZR1) and BRI1-EMS-SUPPRESSOR 1 (BES1), key TFs in transducing BR signals, interacting with PIF4 to regulate a common set of target genes (Oh *et al*., 2012; Martínez *et al*., 2018). Transcriptional activated genes by PIFs include auxin biosynthetic genes which drive enhanced cell elongation rates triggered by temperature and other stimulus (Koini *et al*., 2009; Lucyshyn and Wigge, 2009; Franklin *et al*., 2011). HY5 and PIFs play central roles in a dichotomy controlling hypocotyl cell elongation in which several signalling pathways converge and where signals can be amplified (Pfeiffer *et al*., 2014; Cañibano *et al*., 2021).

Due to its developmental relevance and its utility as a straightforward model for investigating shoot cell elongation, hypocotyl elongation has proven to be instrumental in studying molecular interactions among components of various signalling pathways. This model not only contributes to our comprehension of molecular regulators in response to light but also to other stimuli such as temperature or stress responses and has advanced our understanding of molecular networks within hormone signalling pathways (Wit *et al*., 2016). Auxins are considered to be the principal driver in hypocotyl elongation responses with recent findings supporting the “acid growth” theory (Kutschera, 1994; Arsuffi and Braybrook, 2018; Li *et al*., 2022). Auxin induces the extrusion of protons to the apoplast leading to its acidification which activates expansins at the cell wall. This weakens the wall and induces growth (McQueen-Mason *et al*.,1992). In this process the activation/inactivation of the plasma membrane (PM) H^+^-ATPases by, phosphorylation/dephosphorylation is key and sufficient to promote cell elongation (Takahashi *et al*., 2012; Fendrych *et al*., 2016; Du *et al*., 2020). Auxins induce the expression of SMALL AUXIN UP-RNA (SAUR) from which SAUR19 (and others) activates PM H^+^-ATPase by directly binding and inactivate PROTEIN PHOSPHATASES TYPE 2C (PP2C)-D subfamily of type 2C (PP2C.D) phosphatases, which can render PM H^+^- ATPases inactive by dephosphorylation (Spartz *et al*., 2014). Moreover, auxins can rapidly bind to TRANSMEMBRANE KINASE (TMK) proteins that can directly bind and phosphorylate PM H^+^-ATPases in its penultimate Threonine (Thr^947^) residue promoting hypocotyl elongation (Dai *et al*., 2013; Li *et al*., 2021; Lin *et al*., 2021). Despite auxins being essential to promote hypocotyl elongation and PIFs inducing auxin accumulation in the dark (Franklin *et al*., 2011; Hornitschek *et al*., 2012; Sun *et al*., 2012; Gao *et al*., 2022), it has been demonstrated that auxin transport is required for hypocotyl elongation in light-grown but not dark-grown Arabidopsis (Jensen *et al*., 1998). This idea is supported experimentally by the fact that etiolated hypocotyls are insensitive to NPA, an inhibitor of auxin polar transport whereas etiolated ones are NPA-sensitive (Jensen *et al*., 1998).

Considered as a stress signalling hormone, abscisic acid (ABA) regulates many physiologic stresses as seed dormancy, resistance to drought, stomatal closure and cell elongation (Hirayama and Shinozaki, 2007; Cutler *et al*., 2010). ABA signalling starts with ABA binding to PYRABACTIN RESISTANCE 1 (PYR1)/PYR1-LIKE (PYL)/REGULATORY COMPONENTS OF THE ABA RECEPTOR (RCAR) protein family members (PYR/PYL/RCAR) (Ma *et al*., 2009; Nishimura *et al*., 2009, 2010) which strongly increases their affinity for the cladeA of PP2Cs, such as ABA-INSENSITIVE 1 (ABI1), ABI2, HYPERSENSITIVE TO ABA1 (HAB1), and PP2CA (Antoni *et al*., 2011), leading to their inactivation. Thus, SUCROSE NON–FERMENTING-1–RELATED PROTEIN KINASE 2 (SnRK2s), that act as positive regulators in the ABA signal cascade, i.e., SnRK2.2, SnRK2.3, and OPEN STOMATA 1 (OST1)/SnRK2.6 (Ma *et al*., 2009; Park *et al*., 2009) are released to phosphorylate downstream targets, such as ABA INSENSITIVE 5 (ABI5) or ABA RESPONSIVE TRANSCRIPTION FACTORS (ABF)/ ABRE-binding (AREB) to activate ABA response element (ABRE)–driven gene expression (Uno *et al*., 2000; Fujii *et al*., 2007). SnRKs also activate various types of water and ion channels that determine turgor of guard cells and stomatal aperture thus inducing stress responsive genes concomitantly with channel activation (Nakashima *et al*., 2013; Munemasa *et al*., 2015; Miao *et al*., 2021).

In addition, it has been described a dual role for ABA depending on the concentration. At low concentrations ABA is required for growth and development whereas as high concentrations ABA signals as a stress (Rowe *et al*., 2016; Li *et al*., 2017).

Regarding the control of hypocotyl elongation two different roles have been proposed for ABA. Endogenous ABA is needed for growth and hypocotyl elongation, as shown by the short hypocotyl phenotype of the mutants deficient in ABA accumulation in Tomato, with *sitiens* (*sit*) and *notabilis* (*not*), and in Arabidopsis with *aba1* (Barrero *et al*., 2008; Humplík *et al*., 2015). On the contrary, exogenously applied ABA has long been reported to reduce etiolated hypocotyl elongation, initially in squash hypocotyl segments (Hayashi *et al*., 2014). In Arabidopsis, ABA represses hypocotyl elongation by reducing PM H^+^-ATPase Thr^947^ phosphorylation in an ABI1-dependent manner (Hayashi *et al*., 2014). Recently, an ABI1 direct interaction with PM H^+^-ATPase has been demonstrated with direct consequences in root growth (Miao *et al*., 2021). Moreover, ABA affects the expression of *KAT1* an inward-rectifying K+ channel in etiolated seedlings (Hayashi *et al*., 2014). In fact, it has been recently shown that the *KAT1* promoter is a direct genomic target of PIF3 protein. PIF3 induces KAT1 accumulation at night which upon activation by blue light in the morning leads to the K^+^ intake driving stomata opening (Rovira *et al*., 2023). Additionally, in an independent study, Lorrai *et al*. (2018), show that ABA negatively controls hypocotyl elongation either in etiolated or de-etiolated hypocotyls by acting on GA metabolic genes, inducing the accumulation of the DELLA proteins GAI and RGA, affecting GA signalling, and repressing auxin biosynthetic genes.

Here, we show that, dark-grown etiolated Arabidopsis hypocotyls are insensitive to ABA whereas roots respond to increasing ABA concentrations. The existence of a constitutively active light pathway in the dark, as in *pifq* or in *cop1* mutants, is sufficient for hypocotyl responsiveness to ABA. Thus, the hypocotyl growth inhibition by ABA requires an active light signalling pathway.

## Results

### Etiolated hypocotyls are insensitive to exogenous ABA

The interplay between ABA and light signalling at the level of hypocotyl elongation has been a subject of limited studies. With the aim of accessing the role of exogenous ABA application in hypocotyl elongation we developed a system were plants could be easily transferred to ABA-containing medium after germination. In this system, seedlings germinated on a nylon mesh, under dark or light conditions, could be rapidly bulk-transferred to a new plate after root and hypocotyl emergence, thus avoiding the delaying effect of ABA on seed germination (Supplementary Figure 1).

Using this strategy, wild-type (WT) seedlings germinated in the dark or under low white light (10 μmol m^-2^ s^-1^) were transferred to plates containing 10 µM ABA or ethanol (mock) and maintained for 4 or 5 days respectively at the same light conditions. Whereas, light-grown WT seedlings display root and hypocotyl sensitivity to ABA, as can be measured by a reduction in length, the dark-grown seedlings show root sensitivity to ABA but hypocotyls remained unaffected (Figure 1A). Hence, this is, to our knowledge, the first demonstration that etiolated hypocotyls are insensitive to ABA and contrasts with previous results by Hayashi *et al*. (2014) and Lorrai *et al*. (2018). We believe difference resides in the time of seedling exposure to light during the experimental procedure, since in these studies plants were manipulated under dim red light or have been subjected to higher times of exposure to light (24h) at the initial step of seed synchronization, respectively.

**Figure 1.**
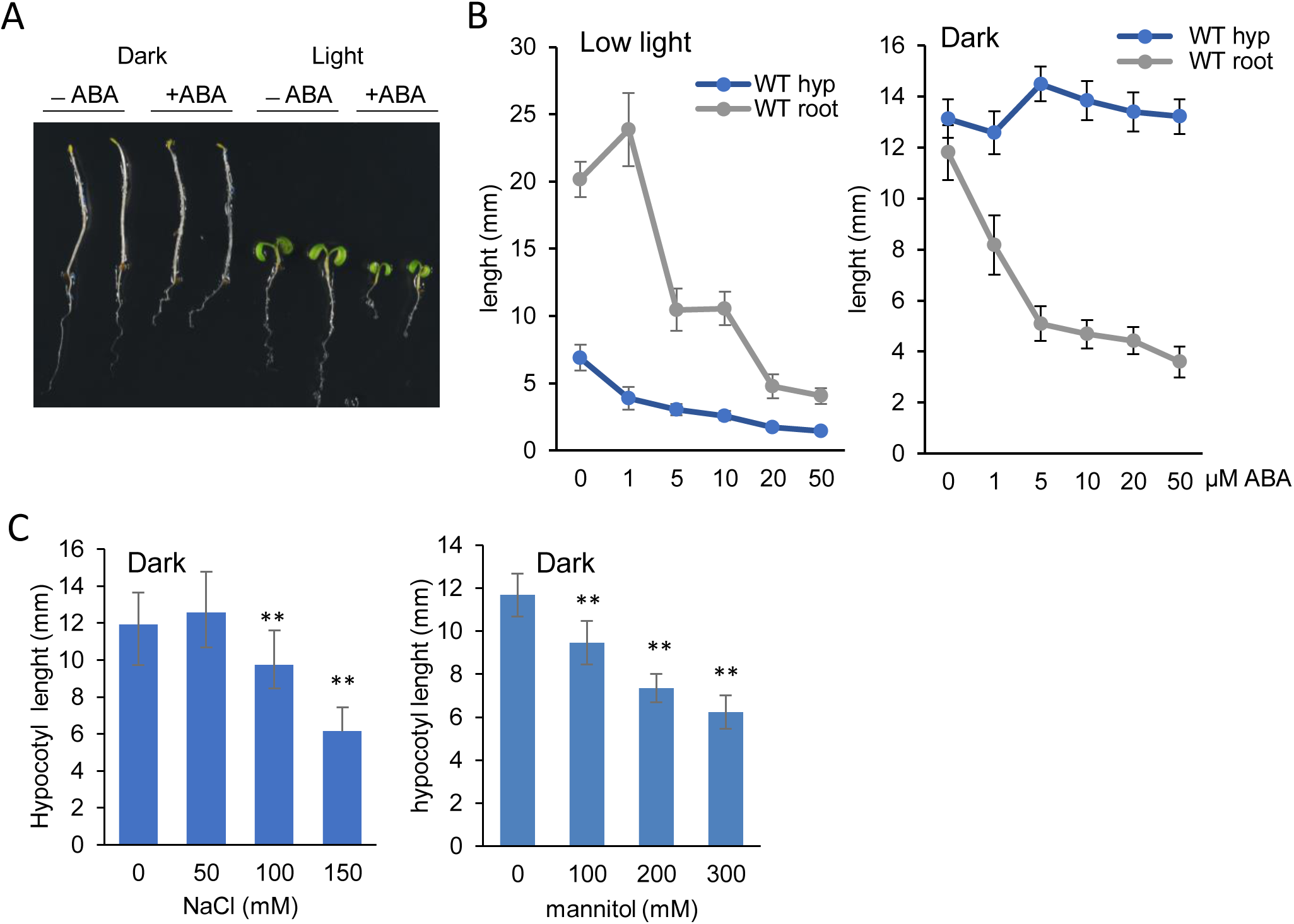
ABA inhibits hypocotyl elongation in the light but not in the dark. **A.** ABA reduces hypocotyl elongation in light grown seedlings but not in etiolated seedlings, whereas roots are sensitive to ABA in both conditions. Seedlings were germinated and kept in media without ABA for 62 hours (dark) or 72 hours (light) and transferred to media with or without ABA (10 µM) for 4 days in the case of dark-grown plants or 5 days for light-grown plants. **B.** Hypocotyls of seedlings grown under low light (10 µm m^-2^ s^-1^) are sensitive to ABA in a dosage-dependent manner. Etiolated hypocotyls are insensitive to ABA up to 50 µM. Roots are progressively responsive to ABA higher concentrations independently of light condition. Lines represent a dosis-responsive curve with the average (n>20) and error bars represent standard deviations. **C.** Etiolated hypocotyls are sensitive to increasing salt and mannitol concentrations. Hypocotyl length of Arabidopsis seedlings grown in the dark for 62 hours that were transferred to plates containing NaCl or mannitol for 4 days. Columns represent average (n>30) and error bars, standard deviation. Asterisks represent significantly different samples (**, p<0.01) regarding the mock control.

When a gradient of ABA concentrations was tested, ranging from 1 to 50 µM, root elongation was progressively impaired in plants growing either under low light or dark conditions (Figure 1B). In a similar way, seedlings germinated under low light showed a progressive reduction in their hypocotyl’s length with increasing ABA concentrations, contrasting with the striking insensitivity of etiolated seedlings up to 50 µM ABA (highest concentration tested). We also observed that in some experiments, in the dark, ABA concentrations of about 5 to 10 µM tend to induce a small increase in hypocotyl length, though this difference was not statistically significant in all the experiments (Figure 1B).

For many physiological responses ABA and salt stress (NaCl) or osmotic stress responses share common regulatory elements. We tested whether WT etiolated hypocotyls display sensitivity to NaCl or mannitol, an osmotic stress inducer. After germination, WT seedlings were transferred to medium containing increasing concentrations of NaCl (up to 150 mM) or mannitol (up to 300 mM), displaying a proportional reduction of hypocotyl elongation after 4 days (Figure 1C). Thus, etiolated hypocotyl sensitivity to salt and mannitol contrasts to the insensitivity to high ABA concentrations, suggesting that salt and osmotic stresses operate by a different path than ABA-induced stress to control hypocotyl elongation.

### Hypocotyl sensitivity to ABA requires an active light signalling pathway

Since the presence of light seems essential for hypocotyl responsiveness to ABA, we tested ABA effect on hypocotyl elongation in mutants where a light signalling pathway is, at least, partially active in the dark, by using *cop1-4* and the *quadruple pif* mutant *pifq* (*pif1 pif3 pif4 pif5*) (Deng *et al*., 1991; Leivar *et al*., 2008). Both mutants showed, when grown in the dark, a reduction of hypocotyl length when seedlings were transferred to plates containing 10 µM ABA (Figure 2A, B). This indicates that an active light signalling pathway, downstream COP1 and PIFs, is necessary for hypocotyl sensitivity to ABA. It is of note that neither *pifq* or *cop1-4* accumulate chlorophyll nor perform photosynthesis in the dark (Deng *et al*., 1991; Leivar *et al*., 2009), suggesting that photosynthetic capacity is not a requisite for ABA sensitivity.

**Figure 2.**
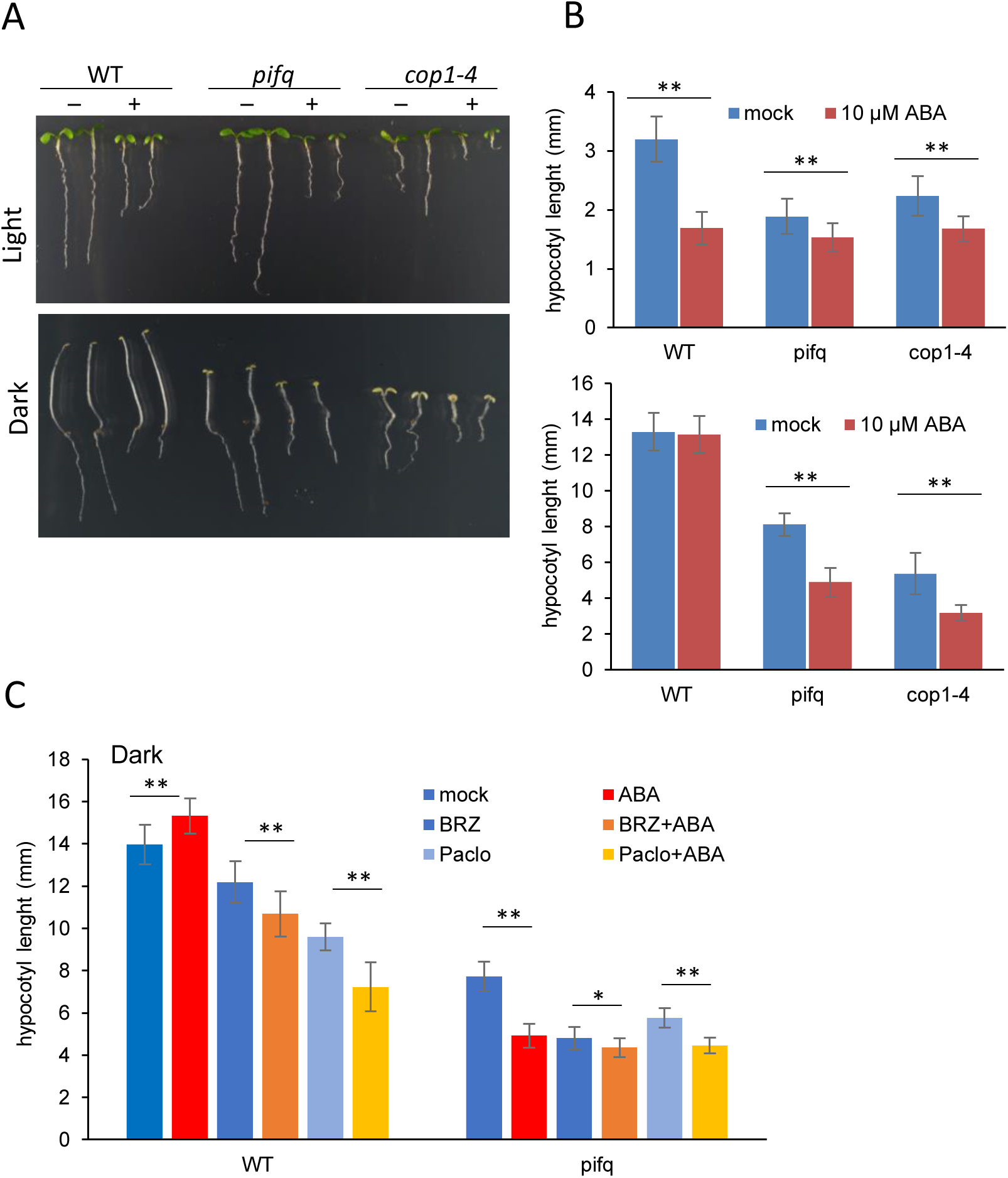
An active light signalling pathway is necessary for hypocotyls responsiveness to ABA. **A.** Hypocotyl elongation of deetiolated *pifq* and *cop1-4* mutants grown in the dark is inhibited by ABA. Seedlings were germinated in media without ABA for 62 hours (dark) or 72 hours (light) and transferred to media with or without ABA (10 µM) for four days for dark grown-plants or five days for light-grown plants. **B.** Hypocotyl length measurements corresponding to the experiments performed in (A). **C.** Hypocotyl length of WT and *pifq* seedlings treated as in (A) but transferred to medium supplied with mock, Brassinazole (BZR) or Paclobutrazol (Paclo) with or without ABA (10 µM). Columns represent average (n>20) and error bars, standard deviation. Asterisks represent significantly different samples (**, p<0.01; *, p<0.05) regarding the non-ABA treated controls.

Additionally, we treated seedlings grown in the dark with Paclobutrazol (Paclo), an inhibitor of GA biosynthesis and with Brassinazole (BRZ) an inhibitor of BR biosynthesis, which by themselves lead to a reduction on hypocotyl elongation (Figure 2C) as expected, since both GAs and BRs are necessary for hypocotyl elongation in the dark (Bouquin *et al*., 2001; Steber and McCourt, 2001; Alabadí *et al*., 2004). When dark-grown seedlings were transferred to plates containing ABA and Paclo, or ABA and BRZ, the hypocotyl length was reduced compared to the controls of plants treated solely with Paclo or BRZ. Thus, the inhibition of the synthesis of GAs or BRs rendered plants sensitive to ABA. Paclo stabilizes the DELLA proteins that repress PIF function (De Lucas *et al*., 2008; Feng *et al*., 2008) whereas PIFs are also required for the BRs responsive BES1 and BZR1 function promoting hypocotyl elongation in the dark (Oh *et al*., 2012; Martínez *et al*., 2018). Altogether, our data support the idea that the PIFs are key players in hypocotyl responsiveness to ABA - in the dark PIFs work as repressors of the ABA-mediated inhibition of hypocotyl elongation.

### Mutants partially blind to light display hypocotyl hypersensitivity to ABA

As darkness confers hypocotyl insensitivity to ABA, we then tested whether photoreceptor mutants that display partial blindness to specific light qualities *phyA* (far-red), *phyB* (red), *cry1*, *cry2*, *cry1cry2* (blue) show some degree of insensitivity to ABA. We found that, upon growing in 10 µM ABA for 6 days, *phyA* and *cry2* mutants, display a reduction of hypocotyl elongation similar to that of WT (Figure 3A). *phyB*, *cry1* and *cry1cry2* display a reduction by more than a half of hypocotyl elongation when treated with ABA suggesting that partial blindness to light does not decrease hypocotyl responsivity to ABA, on the contrary, results in ABA hypersensitivity.

**Figure 3.**
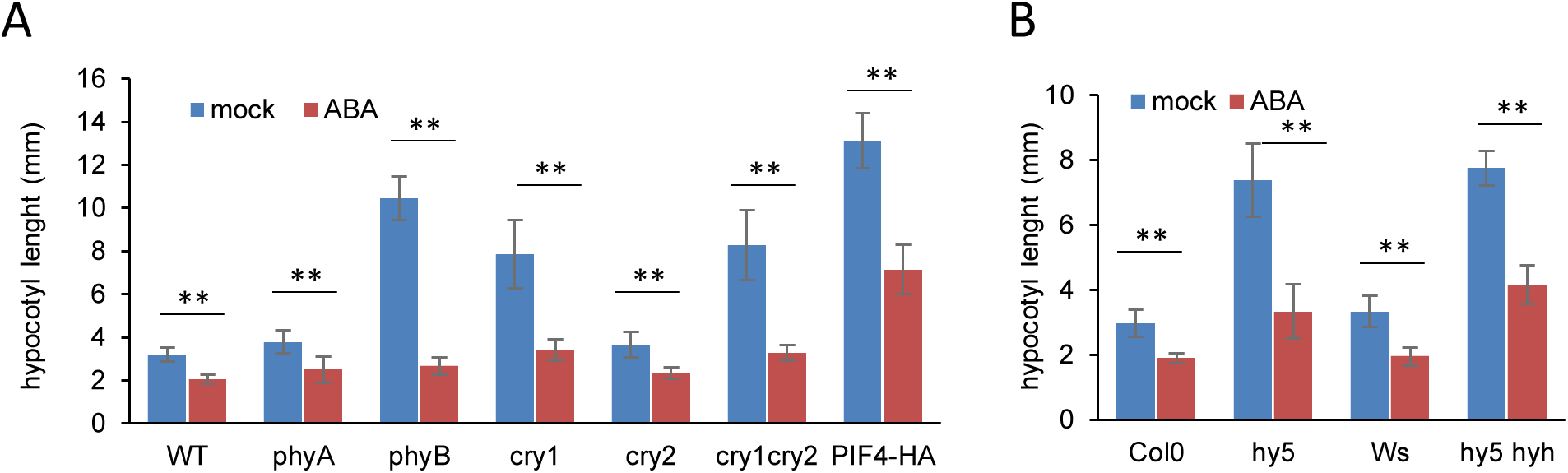
High hypocotyl mutants display enhanced sensitivity to ABA. **A.** Hypocotyl length of WT, photoreceptor mutants and *35S:PIF4-HA* line treated with ABA or mock. **B.** Hypocotyl length of *hy5* and *hy5 hyh* mutants and their respective control backgrounds Columbia (Col) or Wassilewskija (Ws) when treated with ABA. Seedlings were germinated in media without ABA for 72 hours and transferred to media with or without ABA (10 µM) for four days. Columns represent average (n>20) and error bars, standard deviation. Asterisks represent significantly different samples (**, p<0.01) regarding the mock control.

In a similar way, *hy5-215* and *hy5 hyh* mutants that display partial blindness to all light wavelengths and are partially defective in photomorphogenic responses, when transferred to ABA display a stronger reduction of hypocotyl length compared to the WT (Figure 3B). This strong hypersensitivity to ABA might be due to the fact that ABA inhibits hypocotyl growth by reducing cell elongation (Hayashi *et al*., 2014) and, as cells are extremely large in hypocotyl elongated mutants, the action of ABA is more pronounced. In fact, the mutants that already showed low hypocotyls in light as *cop1-4* or *pifq* mutants are suffering a relatively less severe reduction (Figure 2B) which might be because their cells are already small and the effect of ABA is then less pronounced. These results suggest that in light conditions, sensitization to ABA is enabled with consequent hypocotyl size reduction.

Curiously we found that the *35S:PIF4-HA* (De Lucas *et al*., 2008) line which displays high hypocotyls by ectopically accumulating high levels of PIF4 protein is sensitive to ABA in light, in a similar extent to WT, suggesting that PIF4 overaccumulation has no effect in the ABA-mediated inhibition of hypocotyl elongation in light.

### An active ABA signalling pathway is operating in etiolated hypocotyls

It is known that light conditions can regulate the expression of ABA biosynthesis and signalling genes (Huang *et al*., 2019; Yadukrishnan and Datta, 2021). To test if in darkness, the central components of ABA signalling pathway were expressed or if they are, in a general manner induced by light, we tested by qRT-PCR the expression of positive regulators of ABA signalling: ten *PYR/PYL* genes (*PYR1* and *PYL1* to *PYL9*) and *SnRKs2.2* and *2.3* in a transition from dark to light conditions up to 48h. For most of the *PYL* genes tested (exception *PYL7* and *PYL8*) transcripts were induced gradually with light exposure, with *PYL2* and *PYL6* being ten-fold induced at 48 hours, contrasting with the transcript accumulation for the *SnRK2.2* and *SnRK2.3* which showed a slight reduction after 48h of light exposure (Supplementary Figure 2). This data suggest that full expression of the ABA receptors is achieved under light, perhaps due to the onset of tissue specialization, but can hardly justify hypocotyl insensitivity in the dark since there is already a basal level of expression of ABA receptors in the dark.

To test if an active ABA signalling pathway is operating in the dark in WT and *pifq* mutants in a similar way as it is operating in the light, we analysed by qRT-PCR the accumulation of transcripts from ABA reporter genes that are known to be induced by ABA. Transcripts for *ABI1*, *ABI2* and *RD29A* were induced in plants transferred to ABA either in dark or light conditions (Figure 4A). This induction was also observed in *cop1-4* and *pifq* mutants independently of the presence of light (Figure 4A), suggesting that once seedlings are treated with ABA, signal transduction is activated in the dark as in light, triggering stress responses. This signal transduction does not depend on light neither on the presence of PIF1,3,4,5 or COP1 proteins.

**Figure 4.**
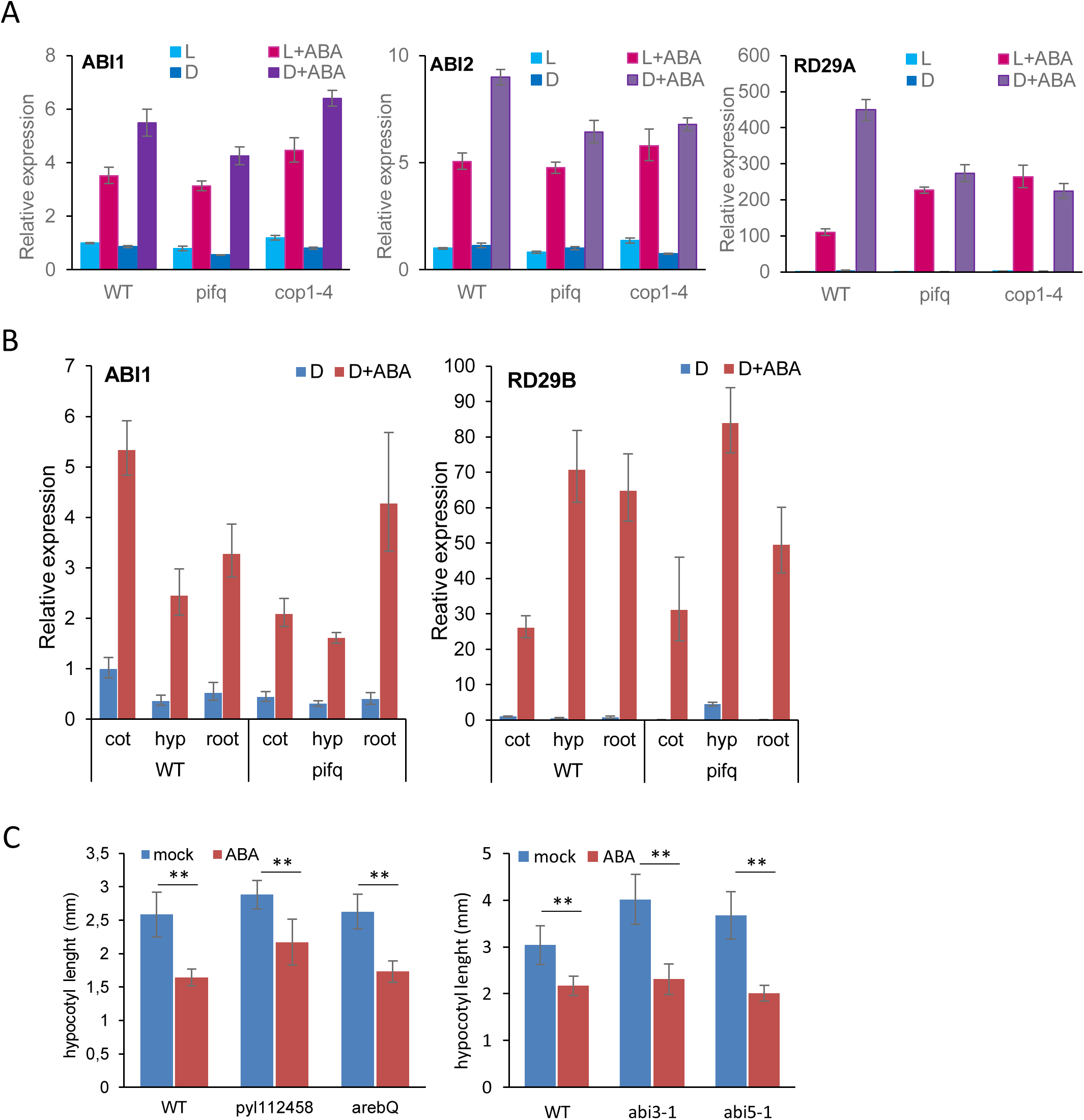
ABA responsive genes are induced by ABA in the dark as in light. **A.** Relative expression of ABA responsive genes obtained by qRT-PCR. *ABI1*, *ABI2* and *RD29A* were induced by ABA treatment either in dark or light conditions in WT and *pifq* or *cop1-4* mutants. **B.** qRT-PCR analysis of gene expression in different tissue samples: cotyledon enriched (cot), hypocotyl (hyp), root enriched (root). *ABI1* and *RD29B* expression is induced by ABA in all the tissues sampled. Columns represent average of four replicates and error bars, standard deviation. **C.** Hypocotyl length of WTs and mutants impaired in ABA signalling grown in low light. Seedlings were germinated in low light for 72 hours and transferred to media with or without ABA (10 µM) for five days. Columns represent average (n>20) and error bars, standard deviation. Asterisks represent significantly different samples (**, p<0.01) regarding the mock control.

We further tested if this ABA induced activation of reporter genes occurs also in the hypocotyl cells unresponsive to ABA in the dark, in a similar way as in other tissues, as roots, that respond to ABA in dark or light. For this, we obtained a cotyledon-enriched fraction (cot), a hypocotyl containing fraction (hyp) and a root enriched fraction (root) from WT and *pifq* plants grown in the dark. Importantly we made sure hypocotyl fraction does not contains tissue from roots or cotyledons. In this plant material the accumulation of *ABI1* and *RD29B* transcripts has been fully induced by ABA in all the tissue fractions isolated in WT and *pifq* mutants, including hypocotyls (Figure 4B). Thus, although etiolated hypocotyls elongation is unaffected by ABA, these hypocotyls display ABA-induced gene expression in a similar way to those of *pifq* mutants which hypocotyl length is reduced by ABA. This indicates that hypocotyl growth inhibition by ABA in the dark is regulated by a different molecular path than that leading to the induction of ABA responsive genes.

We further test if mutants in positive regulators of ABA signalling leading to transcriptional response defects display an impaired hypocotyl responsiveness when grown under low light. For this, we transferred germinated *pyl112458*, *arebQ* (impaired in *ABF1*, *ABF3*, *AREB1*/*ABF2*, *AREB2*/*ABF4*), *abi3-1, abi5-1* and WT seedlings to plates containing 10 µM ABA. Whereas hypocotyl responsiveness to ABA of *arebQ, abi3-1* and *abi5-1* seedlings were similar to that of WT, *pyl112458* sextuple mutant showed higher hypocotyl insensitivity to ABA (Figure 4C), meaning that at least some of the impaired PYR/PYL proteins are important to trigger ABA responsiveness in hypocotyls whereas the TFs ABI5, ABI3, ABF/AREBs are not.

### Transcriptomic analysis of hypocotyl responsiveness to ABA

To identify genes that are important for hypocotyl responsiveness to ABA and whose differential expression levels can justify hypocotyl responsiveness under an active light signalling pathway, we performed an RNA-seq analysis of low-light and dark-grown WT and *pifq* seedlings (later called WT L, WT D, *pifq* L and *pifq* D) transferred to ABA or mock after germination.

We selected the genes being up or downregulated by two-fold by ABA (Log2FC>1 and FDR<0.01) in WT and *pifq* plants grown under dark or light conditions. Since the hypocotyls of WT dark-grown seedlings are insensitive to ABA, and ABA-induced stress responses are active in hypocotyls as in roots, independently of light conditions, we considered all the Differentially Expressed Genes (DEGs) in WT dark-grown plants treated with ABA to be unrelated with hypocotyl elongation. Thus, in order to identify ABA-responsive genes responsible for the reduction in hypocotyl growth under an active light signalling pathway, these genes were subtracted from the ABA-induced and -repressed genes found in *pifq* D, *pifq* L and WT L. To ensure that the resulting lists of genes of interest were flushed more thoroughly of all the ABA-responsive genes in WT D, we applied a less stringent threshold of |Log2FC|>0 to WT D DEGs (Figure 5A). This unveiled that several hundreds of genes respond to ABA specifically in the context of an activated light signalling pathway (i.e. in WT L and *pifq* D or L). Specifically, we found 384 and 709 genes induced and repressed, respectively, by ABA in WT L but not in WT D, and 402 and 668 genes induced and repressed, respectively, by ABA in *pifq* D but not in WT D (Figure 5B).

**Figure 5.**
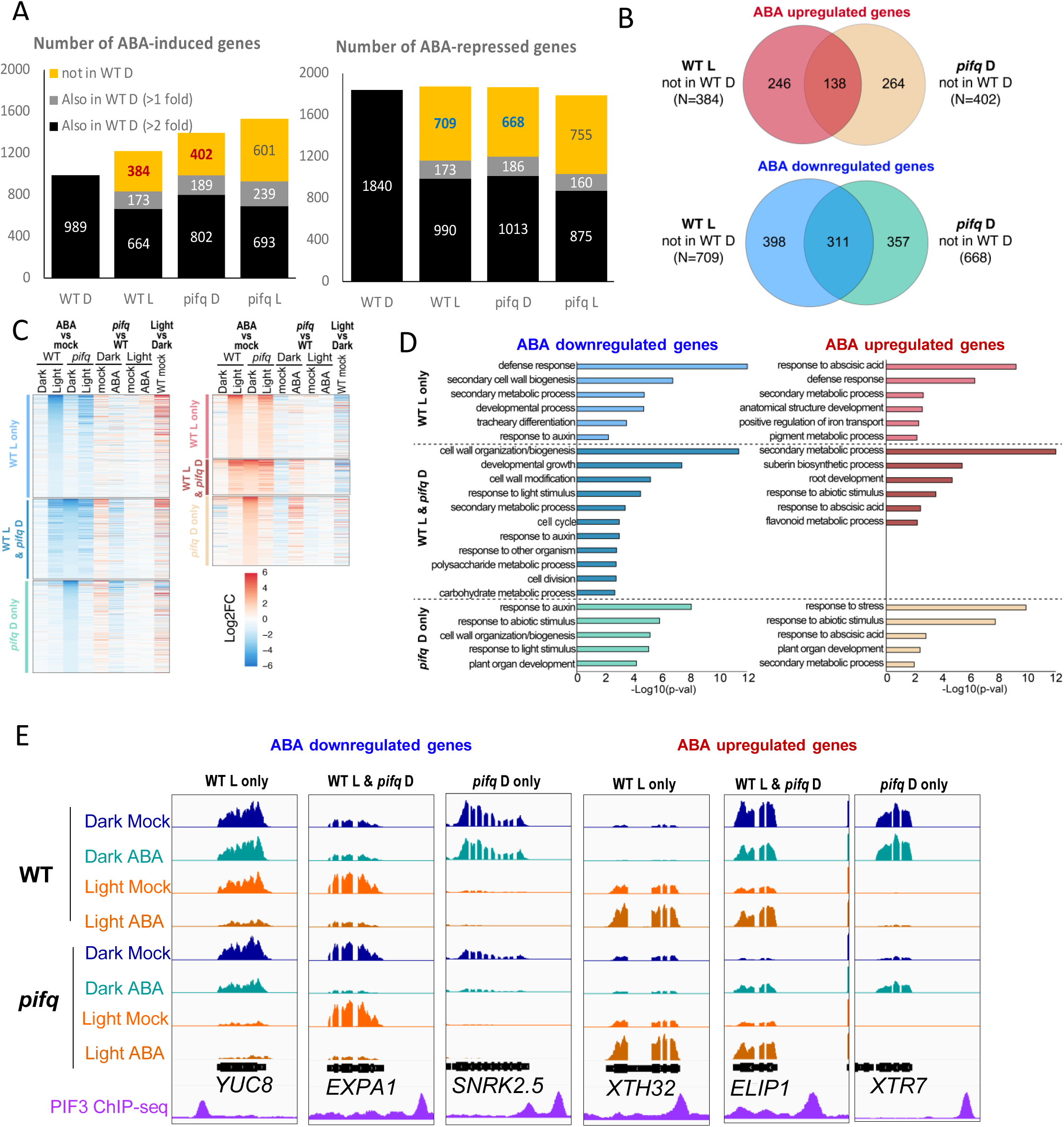
Effect of light and *pifq* mutant background on ABA-induced changes in gene expression. **A.** Number of genes upregulated or downregulated by ABA in wild-type and *pifq* seedlings grown in darkness (D) or light (L) (FDR<0.01 & Log2FC > 1). The number of overlapping genes with WT D are displayed within the WT L, *pifq* D and *pifq* L categories allowing to select the set of genes specifically down- or up-regulated in that categories but not in WT D. **B.** Comparisons of the selected genes according to (A) between WT L and *pifq* D. **C.** Heatmap representing the Log2FC changes in expression in the different pair-wise comparisons in on the 6 gene categories from (B). **D.** GO analysis on the 6 gene categories from (B). **E.** Genome browser view of the RNA-seq data on selected example gene from the 6 gene categories from (B) (average of 3 biological replicates). The profile of PIF3 in WT D seedlings and HY5 in WT L seedlings and the position of G-box motifs have been retrieved from Cañibano *et al*. (2021).

Next, we compared those sets of ABA-regulated genes specifically in WT L or in *pifq* D but not in WT D. This showed that most of these genes were specific to WT L *or pifq* D as only 138 were common between the 384 and 402 ABA upregulated genes and 311 out of the 709 and 668 ABA downregulated genes in WT L and *pifq* D (Figure 5B, Supplementary materials). The genes sufficient for hypocotyl responsiveness to ABA in the *pifq* mutant and which regulation by ABA is blocked by PIFs in the dark are expected to fall in the categories “*pifq Dark only*” or “*WT Light & pifq Dark*” while the genes that might contribute to hypocotyl inhibition by ABA in light through a path independent from PIFs are expected to fall in the “*WT Light only*” category. The core genes that mediate hypocotyl responsiveness whenever a light signalling pathways is activated fall in the “*WT Light & pifq Dark*” category. The heatmap representation of Log2FC of expression from the different pair-wise RNA-seq comparisons within the six gene categories defined in Figure 5B were shows the efficiency of the gene selection process with genes regulated in “WT light only” categories showing no response in WT D nor *pifq* D and genes in the “*pifq* dark only” categories showing no response in WT D nor L (Figure 5C). The response in *pifq* L largely follows the response in WT L, in agreement with the degradation of PIFs in the light. In addition, ABA-downregulated genes in *pifq* D tend to be more expressed in *pifq* D than in WT D and vice versa for ABA-upregulated genes in *pifq* D (Figure 5C; Supplementary materials).

To gain insight into biological processes involved in the response to ABA, we analysed the Gene Ontology (GO) term enrichment of the six categories defined in Figure 5B. For downregulated genes, GO terms related to cell wall organization and biogenesis, response to auxin and developmental processes where significantly enriched for all the three ABA downregulated classes. The GO terms cell division, cell cycle and carbohydrate metabolic process are enriched in the “*WT light & pifq Dark”* category suggesting that these processes are regulated in both contexts. The GO terms response to light is enriched in the classes “*WT light & pifq Dark”* and “*pifq Dark only*”, supporting a crosstalk between PIF-dependent light signalling and ABA response. The GO term defense is prominent in “*WT light only”*, suggesting that defense-related pathways are only activated in the presence of light (Figure 5D).

For upregulated genes, GO terms that are enriched in all the three defined categories are response to ABA or secondary metabolism. Among the enriched GO terms in ABA upregulated genes in “*WT light only*” are defense, iron transport and pigment metabolism. The ABA upregulated genes in “*WT Light & pifq Dark*” are related to suberin biosynthesis (connecting ABA with the development of secondary cell wall), and flavonoid biosynthesis. The genes specifically induced by ABA in the absence of PIFs, upregulated in “*pifq Dark only*” are also related to the response to stress and organ development. Altogether, this analysis shows that ABA regulates genes involved in stress, secondary metabolism, cell wall remodelling, cell division and response to auxin, suggesting these classes are important for ABA responsiveness in light versus dark, likely with many of them being involved in hypocotyl responsiveness to ABA.

### Genes involved in hypocotyl growth or in cell elongation that are responsive to ABA in light or in *pifq*

Since our previous analysis of hypocotyl inhibition by ABA suggested that it is independent of the transcriptional responses driven by ABA TFs (ABI3, ABI5 ABF/AREBs), we looked among our datasets for genes that have been previously linked to hypocotyl elongation either by acting directly at the level of the PM, on the regulation of transporters, channels, that can be connected to the activity of PM H^+^-ATPase, or are involved in auxin synthesis, transport or metabolism, thus having direct consequences on cell elongation. In addition, we considered enzymes involved in cell wall modifications as they might work as a consequence and also as drivers of cell elongation (Cosgrove, 2015).

SAUR gene expression is known to be regulated by PIFs and auxins, and a number of them have been associated with hypocotyl elongation (Koini *et al*., 2009; Lucyshyn and Wigge, 2009; Franklin *et al*., 2011; Leivar and Quail, 2011; Chae *et al*., 2012; Spartz *et al*., 2012; Sun *et al*., 2016; Pfeiffer *et al*., 2014; Ren and Gray, 2015; Gao *et al*., 2022). We found eight *SAUR* genes, (*SAURs 9, 10, 11, 16, 17, 63, 34 and 51*) to be downregulated by ABA specifically in *pifq* D, six SAUR genes (*SAURs 1, 3, 5, 12, 28, 62)* being repressed by ABA in *pifq* D and WT L and nine SAUR genes, (*SAURs 20, 21, 22, 25, 26, 29, 54, 56, 68*) being downregulated by ABA exclusively in WT L. In addition, five SAURs (*SAURs 32, 35, 59, 71, 72*) are significantly induced by ABA only in WT L. In total, 28 members out of a family of 80 members (Markakis *et al*., 2013) are DEGs (mostly downregulated) upon ABA treatment in light or *pifq* in the dark but not in the WT Dark, correlating with the hypocotyl inhibition caused by ABA. Thus, SAURs might play an important role in ABA-mediated inhibition of hypocotyl elongation.

From the auxin related genes that have been previously involved in hypocotyl elongation we identified genes among all the three categories of ABA downregulated genes: *IAA5* being downregulated in “*pifq Dark only*”, *IAA6/SHY1* being downregulates by ABA in WT L and *pifq* D and *IAA19*, together with genes involved in auxin biosynthesis as *TAA1, YUC3, YUC8* and *YUC9*, suggesting that ABA downregulates auxin signalling and biosynthesis under an active light signalling pathway.

We also found a number of ABC transporters to be downregulated by ABA, some of them being previously related to auxin transport (Santelia *et al*., 2005; Lewis *et al*., 2007; Titapiwatanakun and Murphy, 2009; Kaneda *et al*., 2011; Cho and Cho, 2013; Xu *et al*., 2014). *ABCB19,* a well described auxin transporter is among the ABA downregulated genes specifically regulated by ABA in *pifq* D (Supplementary Figure S3B and Supplementary materials). In addition, three ABC-2 type transporters (*ABCG4, ABCG42, ABCG8*) are downregulated in “*WT Light & pifq Dark*” and seven ABC transporters (*ABCB3, ABCB4, ABCB14, ABCC9, ABCG13, ABCG18, ABCG6*) are specifically downregulated by ABA in “*WT Light*”. Of these, ABCB4 and ABCB14 have been previously related to auxin transport (Santelia *et al*., 2005; Cho and Cho, 2013; Xu *et al*., 2014).

Among the DEGs that might control hypocotyl elongation directly at the level of the membranes, we spotted a high number of ion exchangers, solute transporters or channels (Supplementary materials), that by regulating ion homeostasis can directly or indirectly contribute to the control of cell expansion. In particular, we identified the downregulation of *KAT1* and *KAT2* in the PIF-dependent categories (respectively “*pifq D only*” and “*WT Light & pifq Dark*”) as genes for which ABA regulation in the darkness is blocked by PIFs (Supplementary Figure S3B and Supplementary materials). Both genes have been described as inward rectifying channels to compensate for the PM H^+^-ATPase activity (Pilot *et al*., 2001, 2003; Szyroki *et al*., 2001; Véry and Sentenac, 2003; Pandey *et al*., 2007; Lebaudy *et al*., 2008). In addition, the gene encoding AKT2, a photosynthate- and light-dependent inward rectifying potassium channel, gated by Ca^2+^ that can co-assemble with KAT1 (Sharma *et al*., 2013) is upregulated by ABA.

Among the genes associated with cell elongation, we found several belonging to cell wall remodelling families to be DEGs in response to ABA. Among the downregulated, we found seven expansins (*EXPA1, EXPA5, EXPA6, EXPA8, EXPA10, EXPA15* and *EXPB1*), two extensins (*EXT23, EXT33*) and three XTHs (*XTH4, XTH8, XTH19*) and 18 genes involved in pectin modifications. Among the upregulated, *EXPA7, EXPA18, EXT7*, eight XTHs (*XTH9, XTH13, XTH14, XTH16, XTH15/XTR7, XTH22, XTH23, XTH32*) and eight genes involved in pectin modifications (Supplementary Figure S3B and Supplementary materials). These DEGs highlight the tremendous changes triggered by ABA at the level of cell wall reorganization that are essential to change the velocity of cell expansion.

### Direct PIF3 and PIF4 targets are DEGs by ABA

We further inspect the ABA DEGs for the presence of direct PIF3 or PIF4 targets. We identified 74 direct PIF3 targets and 243 PIF4 targets among the ABA-downregulated genes, and 50 PIF3 targets and 144 PIF4 targets among the ABA-upregulated genes. Among the total number of DEGs, PIF3 and PIF4 targets are homogeneously distributed across the six categories and are significantly enriched regarding the expectancy for the total genome (Supplementary Figure S3C). Among the direct PIF4 targets, several genes related with auxin synthesis, signalling and transport, as *YUC8, TAA1, IAA19, IAA29*, several *SAURs, ABCB19* and *AXR3*. The *KAT1* channel, *ABA-INSENSITIVE PROTEIN KINASE 1 (AIK1)* or the *RECEPTOR-LIKE PROTEIN KINASE 1 (RPK1*) are among other target genes previously associated with ABA responses (Supplementary Figure S3B; Supplementary materials). Among the direct PIF3 targets are several *SAURs, IAA19, YUC8 EXPA1, SnRK2.5* and several light and cell wall related genes (Figure 5E and Supplementary materials).

### The inhibition of auxin transport by NPA renders light-grown seedlings’ hypocotyls insensitive to ABA

As shown by the RNA-seq analysis, among the genes that are DEGs by ABA under an active light signalling (in WT L or *pifq* D) and that remain unchanged in WT D, several genes related to auxin signalling, biosynthesis and transport were identified. Despite recent findings supporting a role for auxin signalling on cell wall acidification and thus the acid growth theory (Fendrych *et al*., 2016; Li *et al*., 2021; Lin *et al*., 2021) it has been reported long ago that auxin transport (blocked by N-1-naphthylphthalamic acid; NPA) is required for hypocotyl elongation in light but not in the dark (Jensen *et al*.,1998). Since we also identified a differential effect on hypocotyl inhibition by ABA in plants grown under light or dark conditions, we looked if NPA could affect responsiveness to ABA.

Thus, we tested if blocking auxin transport with NPA affects hypocotyl responsiveness to ABA. In the dark, NPA treatment did not affect WT nor *pifq* hypocotyl elongation, neither affect insensitivity of etiolated plants to ABA. Moreover, in the dark, NPA does not affect *pifq* sensitivity to ABA, suggesting that NPA-inhibited auxin transport does not play a role in ABA sensitivity in the dark, even in the absence of PIFs (Figure 6A).

**Figure 6.**
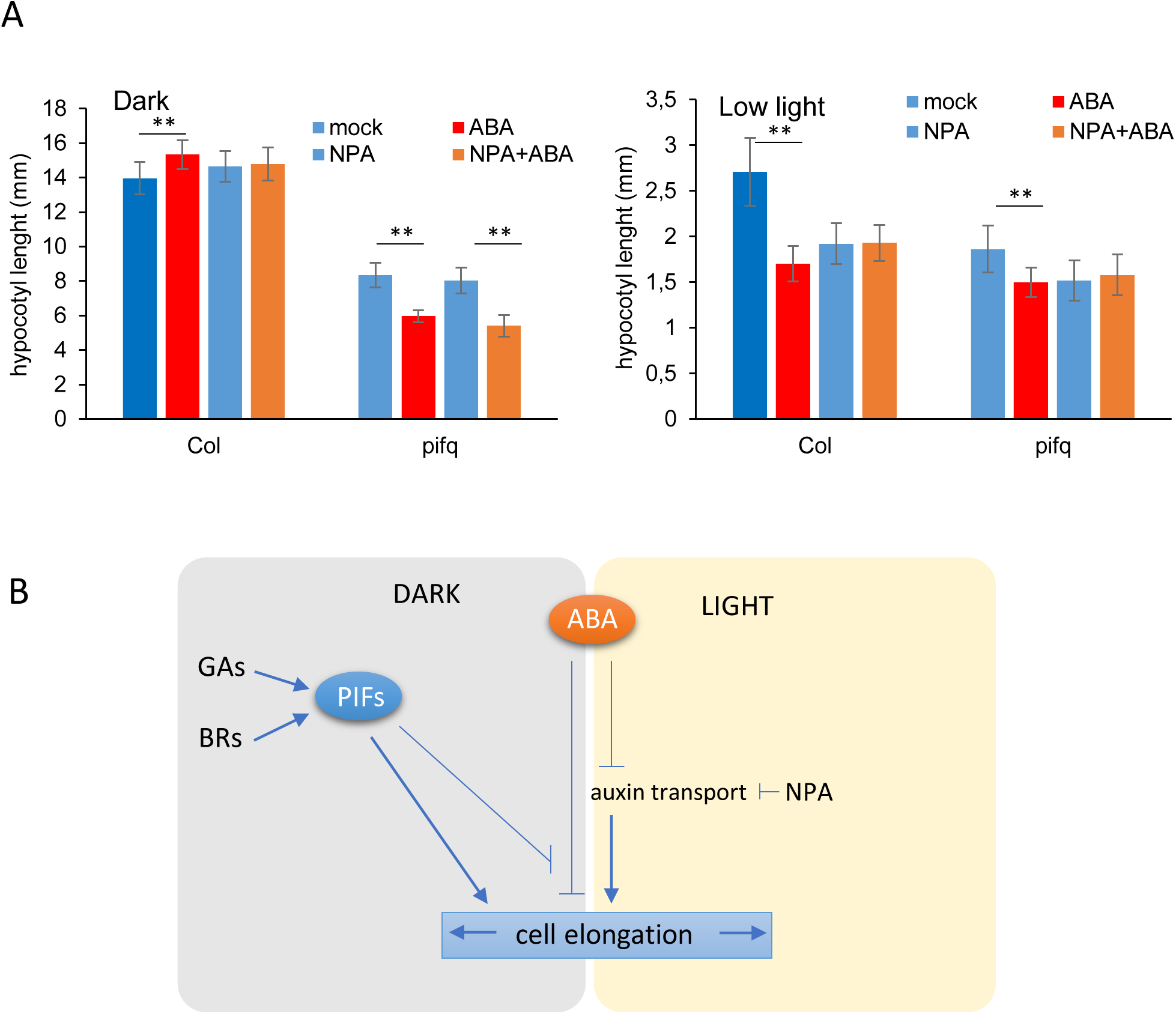
ABA repression of hypocotyl elongation in light requires auxin transport. **A.** Hypocotyl length of WT and *pifq* seedlings grown under dark or low light conditions. Seedlings were germinated in media without ABA for 62 hours (dark) or 72 hours (light) and transferred to media supplemented with ABA (10 µM) and or NPA (0.5 µM) for four (dark) or five days (light). Columns represent average (n>20) and error bars, standard deviation. Asterisks represent significantly different samples (**, p <0.01) regarding the mock control. **B.** Proposed model for ABA inhibition of hypocotyl elongation. ABA repression of hypocotyl elongation in light requires an active auxin transport. In the dark hypocotyls are insensitive to ABA. This insensitizaton is mediated by PIFs that act directly by activating the transcription of genes involved in cell elongation, which in dark is known to be independent on auxin transport (Jensen *et al*.,1998).

In WT light grown seedlings, the presence of NPA resulted in shorter hypocotyls. Furthermore, in the presence of NPA, ABA had no further effect on hypocotyl growth inhibition. Similar results were obtained in *pifq* mutants (Figure 6A). These results show that ABA inhibition of hypocotyl elongation in the light is dependent on a functional auxin transport, which works independently of PIFs. This suggests the existence of an ABA-dependent inhibition of hypocotyl in the light which is either mediated by auxin transport or converge in one or in a set of common molecular targets that requires auxin transport for activation in the light. This path differs from the one that in the dark is fully inhibited by PIFs. It might also be that, in the dark, PIF regulation of cell expansion is acting more downstream than auxin transport, co-regulating directly the elongation capacity at the PM, and thus, this regulation cannot be interfered by auxin transport or ABA (Figure 6B).

## Discussion

Being driven by cell elongation, hypocotyl growth has been used as a fast responsive and hormone sensitive system to study molecular interactions subjacent to cell elongation. The output of hypocotyl elongation rates is the result of the integration of exogenous stimuli into central pathways that directly control growth and that allow the adaptation to changing environmental conditions presenting different stress cues. In nature, for some species, hypocotyl elongation, at early stages of seed development, represents the capacity to emerge from soil and perceive light ensuring the establishment as autotrophic self-sustained organism. Once this establishment occurs, at the initial developmental stages, hypocotyl elongation is crucial for early plant responsiveness to environmental light conditions as light cues, temperature, hypoxia or drought (Boron and Vissenberg, 2014; Krahmer and Fankhauser, 2023).

Regarded as an abiotic stress signalling hormone, ABA’s effect on growth stunting is well known. However, the number of studies on ABA effect on controlling hypocotyl elongation is limited (Wakabayashi *et al*.,1989; Hayashi *et al*., 2014; Lorrai *et al*., 2018). In this study, we report that exposure to ABA reduces shoot size and reduces hypocotyl elongation in light-grown plants. Moreover, we found that etiolated hypocotyls that have not been exposed to light are insensitive to ABA, whereas roots are sensitive to ABA independently of light conditions. As ABA concentration can impact plant development and growth in different ways, with low ABA concentrations promoting growth and high ABA repressing it (Takahashi, 1972; Sharp *et al*., 2000), we tested if high concentrations of ABA can inhibit hypocotyl growth, a phenotype which we could not observe despite 50 µM ABA were added to the media. Our results clearly differ from those obtained by other groups which could detect ABA inhibition of etiolated hypocotyls (Wakabayashi *et al*., 1989; Hayashi *et al*., 2014; Lorrai *et al*., 2018). This might be due to the different experimental setups developed, including the fact that in these previous works plants were manipulated under dim red light (Hayashi *et al*., 2014) or have been subjected to higher times of exposure to light at the initial step of seed synchronization (Lorrai *et al*., 2018).

In fact, the finding that deetiolated mutants (*pifq* and *cop1-4*) grown in the dark display hypocotyl sensitivity to ABA suggests that an active light signalling pathway is necessary for hypocotyl sensitization to ABA, reproducing ABA-sensitivity of light-grown plants (Figure 2A and B).

The finding that the induction of ABA-responsive genes in etiolated hypocotyl is similar to the induction in light, suggests that the ABA-stress responsive pathway is intact in etiolated hypocotyls despite the insensitization to ABA for hypocotyl growth. It also discards the possibility of ABA transport from roots to shoots being impaired in the dark. Thus, ABA acts through divergent paths controlling stress responses and growth, the latest being inhibited by PIFs in the dark. This is in line with the finding that mutants for positive regulators of ABA signalling pathways necessary for transcriptional reprogramming in response to ABA, such as SnRKs and the transcription factors ABI5, ABI3 and ABF/AREBs, show hypocotyl sensitivity to ABA in low light conditions similar to that of WT plants.

It has been reported that salt can impair hypocotyl elongation induced by shade, through the action of the ABA responsive ABF/AREB quartet of transcription factors (Hayes *et al*., 2019). We showed that etiolated hypocotyls are sensitive to salt (Figure 1C), showing strong growth reductions and suggesting that salt regulates hypocotyl elongation by a mechanism that differs from that of exogenous ABA application and therefore probably ABA-independent. In accordance, the *arebQ* mutant involved in salt responsiveness (Hayes *et al*., 2019), displays hypocotyl sensitivity to ABA in low light conditions similar to that of WT (Figure 4C), suggesting that ABA impacts hypocotyl elongation by an alternative molecular path.

The transcriptomic analysis of WT and *pifq* plants treated with ABA in the dark allowed the identification of genes being specifically up or downregulated by ABA in the *pifq* mutant, in the WT in light, or both, but whose deregulation is prevented in WT plants in the dark, where PIFs are active. Among the downregulated genes, we found numerous genes encoding proteins that might play a role in controlling cell elongation directly at the plasma membrane. Among these, there are 23 SAUR proteins, including SAUR9 which can directly interact and inactivate members of the D-clade PP2Cs (Spartz *et al*., 2014) and SAUR63 that can directly promote growth at the PM, by antagonizing PP2C.D5 phosphatase and increasing PM H^+^-ATPase activity (Nagpal *et al*., 2022).

The activation of PM H^+^-ATPase has been shown to be sufficient to trigger growth (Fendrych *et al*., 2016) and the regulation of the phosphorylation of some of its residues is key in the control of its activation (Falhof *et al*., 2016; Miao *et al*., 2021). In fact, ABA has been shown to suppress hypocotyl elongation, in part by promoting the dephosphorylation of PM H^+^-ATPase (Thr^947^ residue), in which the PP2C-type phosphatase ABI1 plays a role (Hayashi *et al*., 2014). Recently it was shown that ABI1 can directly bind to PM H^+^-ATPases and inhibit its activity by promoting the dephosphorylation of Thr^947^, controlling root elongation (Miao *et al*., 2021). The repression of *SAUR* expression by ABA can be an additional mechanism of restraining hypocotyl cell elongation by keeping PM H^+^-ATPases in a sustained inactive state. Since *SAUR*s are induced by auxins and are direct targets of PIFs (Zhang *et al*., 2013; Pfeiffer *et al*., 2014; Sun *et al*., 2016), in the dark SAURs are being constantly induced. When SAUR proteins are present, the Thr^947^ of H^+^-ATPase remain phosphorylated and the pump is active (Spartz *et al*., 2014) promoting fast cell elongation. The presence of PIFs in WT D could counteract ABA-driven downregulation of SAURs, a mechanism to directly prevent unwanted inhibition of cell elongation by ABA in dark-grown seedlings.

Among the genes downregulated by ABA in *pifq* that may play a role in cell elongation at the PM we found the potassium channels KAT1 and KAT2 which work as an inward rectifying K^+^ channels for PM H^+^-ATPase activity and are involved in auxin-induced hypocotyl elongation of etiolated seedlings (Takahashi *et al*, 2012). In fact, KAT1 has been shown to be a direct PIF3 target, playing a role on the PIF control of stomata opening at the onset of day periods (Rovira *et al*., 2023). Moreover, we identified AKT2, another inward rectifying channel, regulated by Ca^2+^ that can associate with KAT1 (Ivashikina *et al*., 2005), to be upregulated by ABA, suggesting the control of K^+^ channels, might be a direct path by which ABA reduces hypocotyl cell elongation.

Concomitantly to these, a number of genes encoding for cell wall modification enzymes as EXPs, XTHs, and Pectin melthylesterases (PMEs) are being deregulated by ABA in *pifq* mutant, promoting cell wall rearrangements according to changes in elongation rates. Thus, by activating a number of genes directly interfering with PM H^+^-ATPase function, and cell wall remodelling, PIFs master-regulate cell elongation in the dark, insensitizing ABA signals that could potentially inhibit cell elongation.

Contrary to what could be expected, hypophotomorphogenic mutants affected in the light-signaling pathway that display elongated hypocotyls in the light *hy5*, *hy5 hyh*, *phyB*, *cry1* and *cry1 cry2* display enhanced sensitivity to ABA. This suggest that upon light perception different mechanisms by which ABA can reduce hypocotyl elongation might be activated. These do not depend on PIF repression, as can be inferred by the hypocotyl inhibition of the *35S:PIF4-HA* by ABA, in a similar extent to that of the WT. It might be, that in light conditions, when HY5 is impaired (*hy5* mutant) ABA responsive TFs could gain a prominent role promoting growth reduction in response to ABA, as some of these factors share targets with HY5 and bind also to G-boxes (Cañibano *et al*., 2021; Song *et al*., 2016). Actually, the overexpression of ABI5 restores ABA sensitivity in *hy5* and results in enhanced light responses and shorter hypocotyls in the wild type (Chen *et al*., 2008).

Additionally, the auxin transport inhibitor, NPA, renders light-grown plants insensitive to ABA, suggesting that ABA requires an NPA-repressible auxin transport to inhibit hypocotyl elongation in the light. Since such auxin transport is required for hypocotyl elongation in seedlings grown in the light but not in seedlings grown in the dark (Jensen *et al*., 1998), ABA unresponsiveness is not affected by NPA treatment in the dark. Our transcriptome analysis revealed a number of ABC transporters, some of them previously related to auxin transport (as ABCB4 and ABCB14) to be downregulated by ABA specifically in the light. Curiously the well-studied auxin transport ABCB19 is being downregulated by ABA specifically when PIFs are abscent in the dark (Supplementary Figure S3B). As the direct targets of NPA remain to be unequivocally determined (Teale and Palme, 2018; Geisler, 2021), these results leave space for ambiguous interpretation of auxin transport components downstream ABA signalling. Furthermore, it has been shown that auxin promotes hypocotyl elongation by a PIF4/5-independent pathway during the day (Chapman *et al*., 2012). In our assays, in the *pifq* mutant in the dark, NPA does not reduce hypocotyl elongation neither blocks the effect of ABA on hypocotyl elongation in this mutant, showing that no NPA-dependent auxin transport is needed for the regulation of hypocotyl size in the dark, even in a constitutively photomorphogenic *pifq* genetic background. This supports the idea that NPA blocks the effect of ABA on hypocotyl elongation only in light-grown plants. Thus, the auxin transport (inhibited by NPA) that occurs in the light is independent of PIFs and is, in our experiments necessary for ABA repression of hypocotyl elongation only in the light, advocating for different mechanisms of ABA control of hypocotyl elongation operating in light or dark.

The relation of ABA and auxins for the regulation of cell elongation is complex. The Arabidopsis mutant *aba1* and the tomato mutants *sit* and *not*, deficient in ABA accumulation, show shorter hypocotyls (Barrero *et al*., 2008; Humplík *et al*., 2015) suggesting endogenous ABA is necessary for hypocotyl elongation. ABA has been shown to control local IAA oxidation to regulate IAA homeostasis and hypocotyl elongation in tomato (Lei *et al*., 2023). Moreover, AAO3 mutant, impaired in the last step of ABA biosynthesis is important for the inhibitory effect of auxins in dark-grown hypocotyls elongation, suggesting that ABA might play a role downstream of auxins in this regulation (Emenecker *et al*., 2023). In fact, we detect that ABA can affect the expression of *TAA1* and three *YUCCA* genes which might account for specific relations between ABA and auxins in hypocotyls.

Response to abiotic stresses induced by ABA triggers a transcriptional reprograming that allows plant to adapt to environmental changes, by adapting their morphology and metabolic responses. The control of growth by elongation is part of these responses. Here, we show how inhibition of hypocotyl elongation by ABA occurs in light but not in dark-growing seedlings. These responses might reflect an adaptative mechanism by which the germinating seedling prioritizes reaching light for survival and insensitizes hypocotyl cells to the ABA signal, preventing the reduction of the elongation rate, ensuring to quickly reach the soil surface even under stressful conditions. In this way a sudden shortage of soil water content due to weak water retention, just upon germination would not reduce the possibilities to reach soil surface and perceive light.

## Material and Methods

### Plant Materials

Wild-type and mutant plants used in this study were all in the Columbia-0 (Col-0) ecotypes with the exception of the *hy5 hyh* double mutant (Holm *et al*., 2002) which is in Wassilewskija (WS) ecotype, and *abi3-1* and *abi5-1* mutant (Finkelstein, 1994), in *Landsberg erecta* (*Ler*) ecotype. *phyB* (Reed *et al*., 1993), *phyA* (Somers *et al*., 1998) and *pifq* (Leivar *et al*., 2008) seeds and the *35S:PIF-HA* (De Lucas *et al*., 2008) complemented line were kindly provided by Prof. Salome Prat (CRAG, Barcelona, Spain). The sextuple mutant *pyl112558* (*pyr1 pyl1 pyl2 pyl4 pyl5 pyl8*) (Gonzalez-Guzman *et al*., 2012) was kindly provided by Prof. Pedro L. Rodriguez (IBMCP, Valencia, Spain). *areb1 areb2 abf3 abf1* (*arebQ*) seeds (Yoshida *et al*., 2015) was kindly provided by Prof. Ronald Pierik (Utrecht University, Netherlands). The mutants *cop1-4*, *cry1*, *cry2, cry1 cry2* and *hy5-215* were previously described in McNellis *et al*. (1994), Ahmad and Cashmore (1993), Mockler *et al*., (1999); Somers *et al*. (1998) respectively.

### Plant growth conditions

Arabidopsis seedlings were sterilized with a solution of 75% sodium hypochlorite and 0.1% Tween-20 and stratified at 4°C during 2 days in darkness. Seedlings were seeded on a 350 µm pore nylon mesh (Distribuciones Quimebora S.L., Madrid, Spain) placed over solid Murashige and Skoog (MS) medium with 1% sucrose and 0.6% agar. For experiments in the dark, seeds were exposed to light for 2h to synchronize germination, wrapped in aluminium foil and after 62 hours transferred, on the top of the mesh, to plates containing ethanol (mock) or ABA (Sigma Aldrich) and kept in the dark for four additional days, unless otherwise stated. For experiments under low light, seeds were placed at under fluorescent white light (10 μmol m^-2^ s^-1^) with a 16-h light/8-h dark period and transferred to mock or ABA containing plates after 72 hours and kept under low light for five days, unless otherwise stated. All the plant were grown at 22°C. For the hormone inhibitors, 0.5 uM NPA (Duchefa Biochemie); 0.5 uM BZR (MedChemExpress) and 0.1 uM Paclo (Duchefa Biochemiel) were used). The use of a 350 µm pore mesh reproduced initial results obtained by transferring seedlings to ABA or mock containing media very gently with the help of sterilized forceps.

### Hypocotyl measurements

For hypocotyl measurements light or dark grown plants were disposed on agar plates, photographed and hypocotyls measured using ImageJ software (http://www.imagej.net). At least three biological replicates, each consisting of measurements of at least 20 seedlings grown at different times, were analysed with similar results.

### qRT-PCR

For RT-qPCR assays, 2 µg total RNA was extracted from 7-day old seedlings with the Favorprep Plant Total RNA Purification Mini kit (Favorgen) was used for cDNAs synthesis with using the High-Capacity cDNA Reverse Transcription kit (Applied Biosystems) with DNAse I treatment (Roche). Quantitative PCR was carried out using 5x PyroTaq qPCR mix Plus EvaGreen (CMB Cultek Molecular Bioline) in a QuantStudio5 machine (Applied Biosystems). Transcripts were amplified and results were normalized to *PP2A* transcript levels. Primers used for qRT-PCR are represented in Supplementary Table S1.

### RNA-seq analyses

For RNA-seq, 2 µg total RNA were extracted from three independent biological replicates of WT and *pifq* seedlings, grown under light or dark conditions and transferred to ABA or mock as stated above. Total RNA was extracted with the Favorprep Plant Total RNA Purification Mini kit (Favorgen). Messenger (polyA+) RNAs were purified from total RNA using oligo(dT). Stranded libraries were prepared using DNB-seq (BGI). A 100-bp paired read end sequencing was performed on DNBSEQ-G400 system (BGI). Reads were quality-checked with FASTQC (Andrews, 2010) and mapped on TAIR10 genome assembly of A. thaliana genome using STAR (Dobin et al., 2013) with relevant options “--alignIntronMin 20 --alignIntronMax 5000 --alignMatesGapMax 10000”. Duplicated alignments were removed using samtools markdup -r (Li et al., 2009). Browser tracks were generated using deepTools function bamCoverage with options “--binSize 10 --normalizeUsing RPKM” and visualized using IGV browser (Robinson et al., 2011). Read counts were retrieved using QoRTs (Hartley and Mullikin, 2015) using gene coordinates obtained from file Arabidopsis_thaliana.TAIR10.42.gtf. Differential expression was analyzed with DESeq2 (Love et al., 2014) using default options. Differentially expressed genes for each comparison were selected by adj.p-value < 0.01 and |FC| > 1. Heatmaps and GO enrichment analysis was performed as in Bourbousse *et al*., (2015).

## Acknowledgements

We thank to Salome Prat (CRAG, Barcelona) for the seeds of the light signalling mutants, to Pedro Rodriguez (IBMCP, Sevilla) for sharing seeds of ABA signalling mutants and Scott Hayes and Ronald Pierik (Utrecht University) for sharing seeds of the *arebQ* mutant. Research in SF lab was supported by PID2019-109925GB-I00 funded by MINECO, an Europa Excelencia project EUR2022-134066 funded by Spanish Ministry For Science and Innovation and by a *Beca Leonardo a Investigadores y Creadores Culturales* (IN[21]_CMA_BIO_0136) supported by the BBVA Foundation. This work benefitted from grants from the Agence Nationale de la Recherche projects (httpANR-20-CE13-0028) to CB.

## Supplementary Figures

**Supplementary Figure S1.**
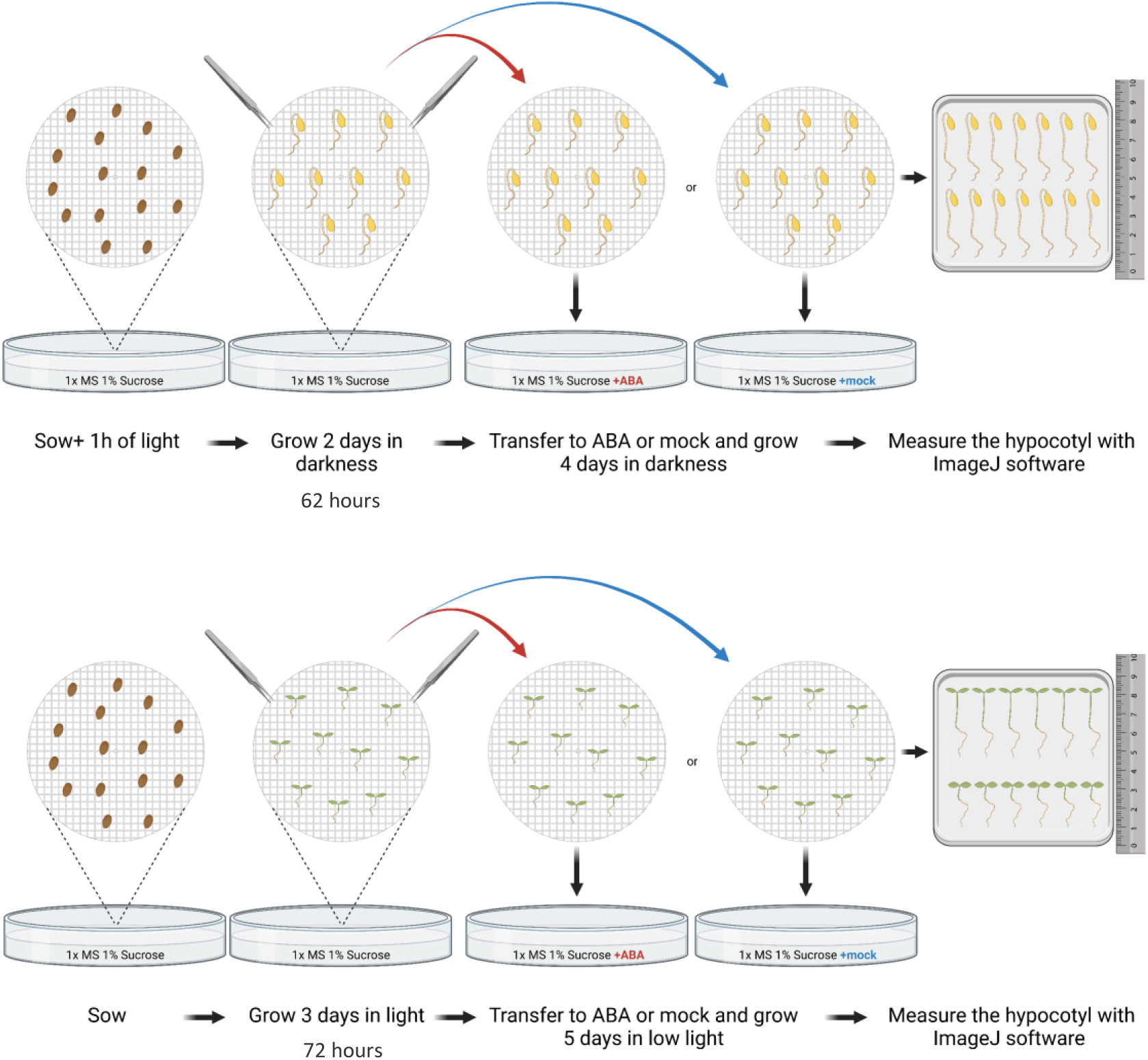
Schematic representation of seedling transfer to ABA or mock containing medium using a mesh. Arabidopsis seedlings were sterilized and sowed in water at 4°c for 48 hours. Then seeds were plated over a 350 µm pore nylon mesh. For dark experiments seeds were kept in light for one hour to synchronize germination and then wrapped in aluminium foil. After 62 hours the mesh containing the germinated seedlings was transferred to a new MS plate containing ABA or mock (ethanol) and kept under dark for 4 days. For light experiments plates were placed under normal light conditions (100 µm m^-2^ s^-1^) and after 72 hours the mesh was transferred to MS medium containing ABA or mock. Then the seedlings were kept under low light conditions (10 µm m^-2^ s^-1^) for 5 days.

**Supplementary Figure S2.**
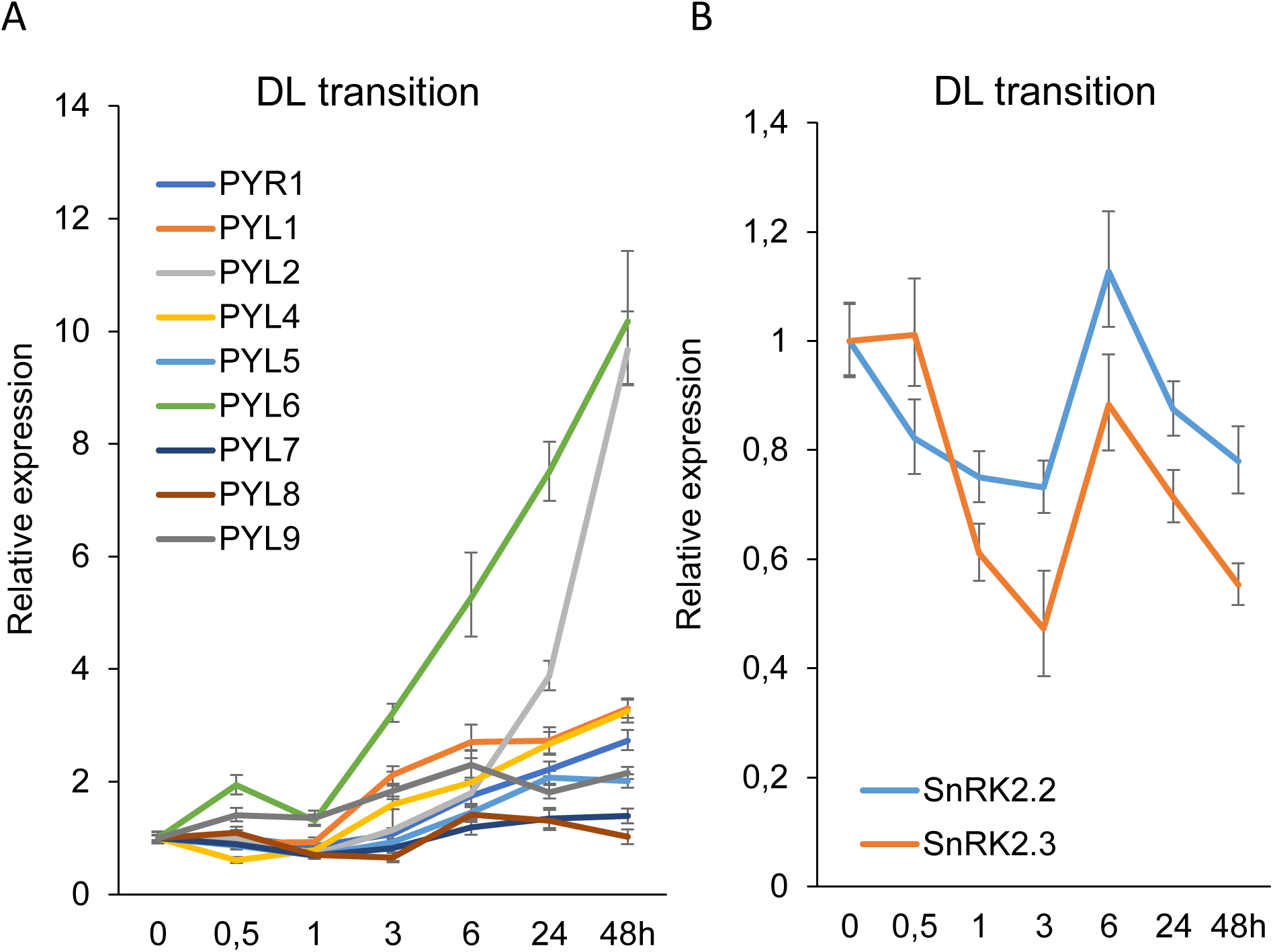
The expression of a number of PYL receptors is induced by light. **A.** qRT-PCR analysis of the *PYR/PYL/RCAR* receptors transcripts during the transition from dark to light over a time course of 48 hours (h). **B.** Transcript accumulation of *SnRK2.2* and *SnRK2.3*. Lines represent transcript accumulation over time and error bars represent the standard deviation of four replicates.

**Supplementary Figure S3.**
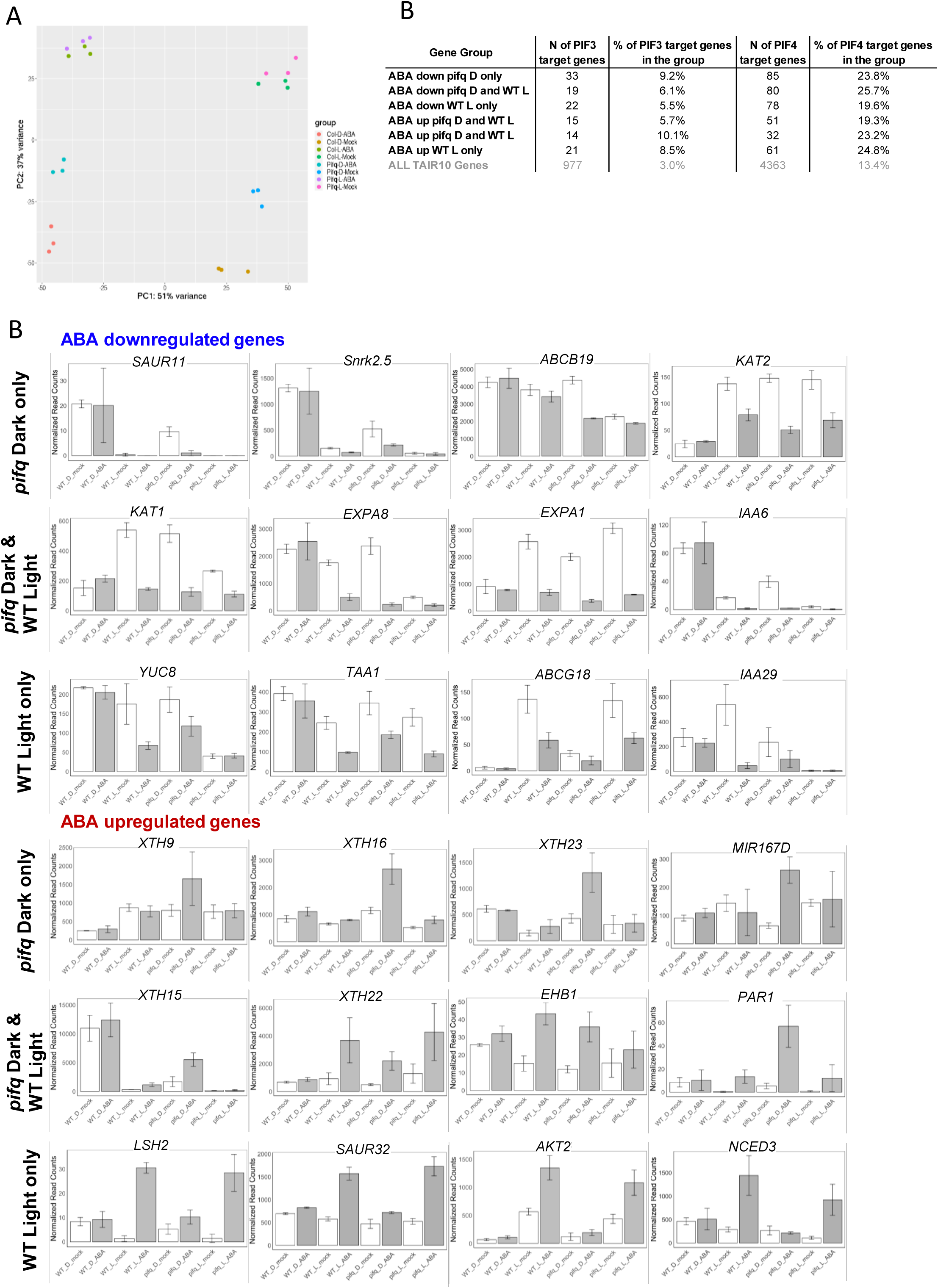
**A.** PCA analysis of the transcriptome dataset. **B.** Normalized expression levels of example genes. Barplots represent the mean and the error bars the standard deviation of the three biological replicates. **C.** Representation of PIF3 target genes within the different ABA-regulated gene groups.

**Supplementary Table S1.**
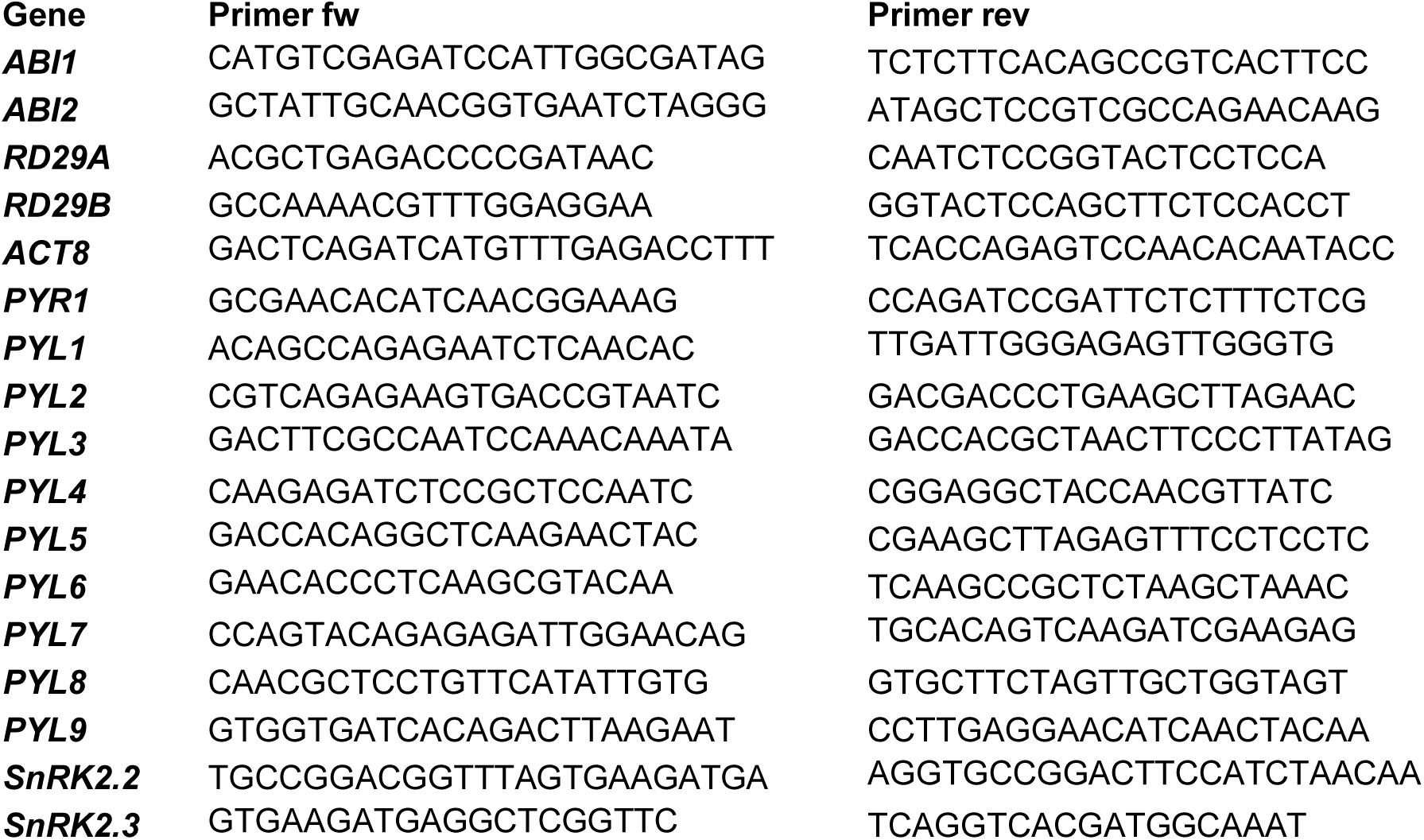
Primers used for qRT-PCR assays.

